# Divergent regulation of STING-mediated innate immune responses by CBP and p300

**DOI:** 10.64898/2026.08.30.745803

**Authors:** Koteswararao Garikapati, Derek Van Booven, Mohammad Faraz Zafeer, Mustafa Tekin, Gaofeng Wang

## Abstract

Dysregulation of cGAS-STING pathway contributes to various disorders, including autoimmunity, infectious diseases and cancer. Transcription factors, particularly IRF3 which presumably recruits CBP/p300 as transcription coactivators, drive innate immune responses to STING activation. Here we show that CBP/p300 inhibitors (p300i) boost, not repress as expected, the cGAMP-induced transcription of innate immune genes. Mechanistically, p300i and a CBP/p300 degrader markedly enhance the cGAMP-induced IRF3 phosphorylation and slightly increase NF-κB activation. Furthermore, cGAMP stimulation increases the presence of p300, not CBP, in the cytoplasm to interrupt interactions between TBK1 and IRF3. Conversely, p300i enhances TBK1-IRF3 interactions, resulting in higher IRF3 phosphorylation. After STING activation, CBP, not p300, is recruited to chromatin along with IRF3 and is coupled with H3K27ac upregulation. At later stages of STING activation, nearly all CBP and p300 are detached from chromatin, which is associated with global histone deacetylation, IRF3 chromatin eviction, and the termination of IRF3-driven transcription programs. p300i robustly enhances IRF3 chromatin engagement which consequently amplifies type 1 interferon responses to STING activation. Together, these findings suggest that CBP, not p300, acts as a transcription coactivator in the nucleus. In contrast, p300, not CBP, is a suppressor of STING signaling in the cytoplasm.

## INTRODUCTION

In mammalian cells, DNA is confined in the nucleus and mitochondria. The presence of DNA fragments in the host cell cytoplasm, derived from either the invaded DNA viruses and bacteria or self-DNA that are damaged and leaked from the nucleus and mitochondria, is detected by the pattern recognition receptor cyclic GMP-AMP synthase (cGAS). cGAS assembles on double stranded DNA (dsDNA) fragments which activate its catalytic activity to synthesize 2′3′-cyclic GMP-AMP (cGAMP), a unique second messenger (1–3). Subsequently, the binding of cGAMP to the stimulator of interferon genes (STING) triggers conformational changes to traffic from the endoplasmic reticulum to the Golgi apparatus to recruit TANK-binding kinase 1 (TBK1). After autophosphorylation and activation, TBK1 phosphorylates the transcription factor interferon regulatory factor 3 (IRF3), specifically at C-terminal regulatory domain, which is an initial step for activating the transcription of innate immune responsive genes, especially type I interferons (IFN) (4–7). TBK1 also coordinates with IκB kinase to activate NF-κB to promote the expression of proinflammatory cytokines such as IL-6 (8, 9). Overall, cGAS-STING pathway constitutes one of the fundamental pathways mediating innate immunity.

The strength and duration of responses elicited by cGAS-STING pathway must be tightly controlled to maintain homeostasis. Overreactions, especially to self-DNA, leads to autoimmune and autoinflammatory disorders, while insufficient responses contribute to the failed host defense against microbial invasion and diseases such as cancer (10, 11). Acetyltransferases CBP/p300 have been thought to synergize with IRF3 to activate the transcription of type 1 IFNs and other innate immune responsive genes by acetylation of histones to increase chromatin accessibility for facilitating transcription factor recruitment and sustaining transcription (12–16). After STING activation, the phosphorylated IRF3 forms dimers to translocate to the nucleus to bind with DNA, especially at interferon-stimulated response elements (ISREs) through the N-terminal DNA binding domain, while presumably recruiting CBP/p300 as transcription coactivators via the C-terminal IRF association domain, thus setting the stage for the transcription of genes such as interferon β (IFNβ).

Intriguingly, there is no previous study that has specifically examined the role of CBP/p300 in cGAS-STING pathway, possibly due to the fact that CBP/p300 have already been established as transcription coactivators. If CBP/p300 only play a transcription coactivator role in cGAS-STING pathway, inhibition of CBP/p300 would be expected to suppress the IRF3-driven gene expression. However, results of our initial experiments unexpectedly show that inhibition of CBP/p300 by selective inhibitors does not inhibit, but robustly enhances, the strength and extends the duration of the transcription of innate immune responsive genes, especially IFNβ and interferon-stimulated genes (ISGs), in response to STING activation. Following our further analyses, we report here that CBP and p300 play distinct roles in cGAS-STING pathway. STING activation increases the presence of p300, not CBP in the cytoplasm, which subsequently interrupts TBK1-IRF3 interactions. Conversely, p300i enhances TBK1-IRF3 interactions and promotes IRF3 activation. At early stages of STING activation, CBP, not p300, is co-recruited to chromatin with IRF3. At later stages of STING activation, nearly all CBP and p300 are detached from chromatin, which is associated with global histone deacetylation and IRF3 chromatin eviction. Overall, CBP serves as a transcription coactivator in the nucleus and p300 acts as a suppressor of STING signaling in the cytoplasm.

## RESULTS

### p300i promotes innate immune gene transcription

We initially investigated p300i SGC-CBP30 on IRF activity, measured by a luciferase assay in RAW-Lucia ISG cells, in which the IRF-inducible luciferase reporter gene was incorporated in the host cell genome. While not directly inhibiting the enzymatic capacity of CBP/p300, SGC-CBP30 blocks its bromodomain to bind to the acetylated lysine residues, thus suppressing further acetylation of substrate proteins. Stimulation with a single dose of cGAMP (2 µM), which can enter the cells via importers (17), robustly induced IRF luciferase activity. Pretreatment with SGC-CBP30 (1 µM) significantly enhanced the cGAMP-induced IRF luciferase activity (Figure 1A). Without SGC-CBP30, the cGAMP-induced luciferase activity peaked at 8h, which was a 3.1-fold increase compared to the baseline. After SGC-CBP30 pretreatment, the cGAMP-induced luciferase activity was risen to 14.8-fold of the baseline and peaked at 16h of cGAMP stimulation. To validate the effects of SGC-CBP30 on IRF activity, we tested another p300i A485 which directly inhibits the catalytic activity of CBP/p300. Consistently, A485 (1 µM) promoted the cGAMP-induced IRF luciferase activity by elevating its peak and by expanding its active window (Figure 1B). Comparatively, the effect of A485 at the same concentration was stronger than that of SGC-CBP30. These results suggest that p300i enhances IRF activity in response to STING activation.

**Figure 1.**
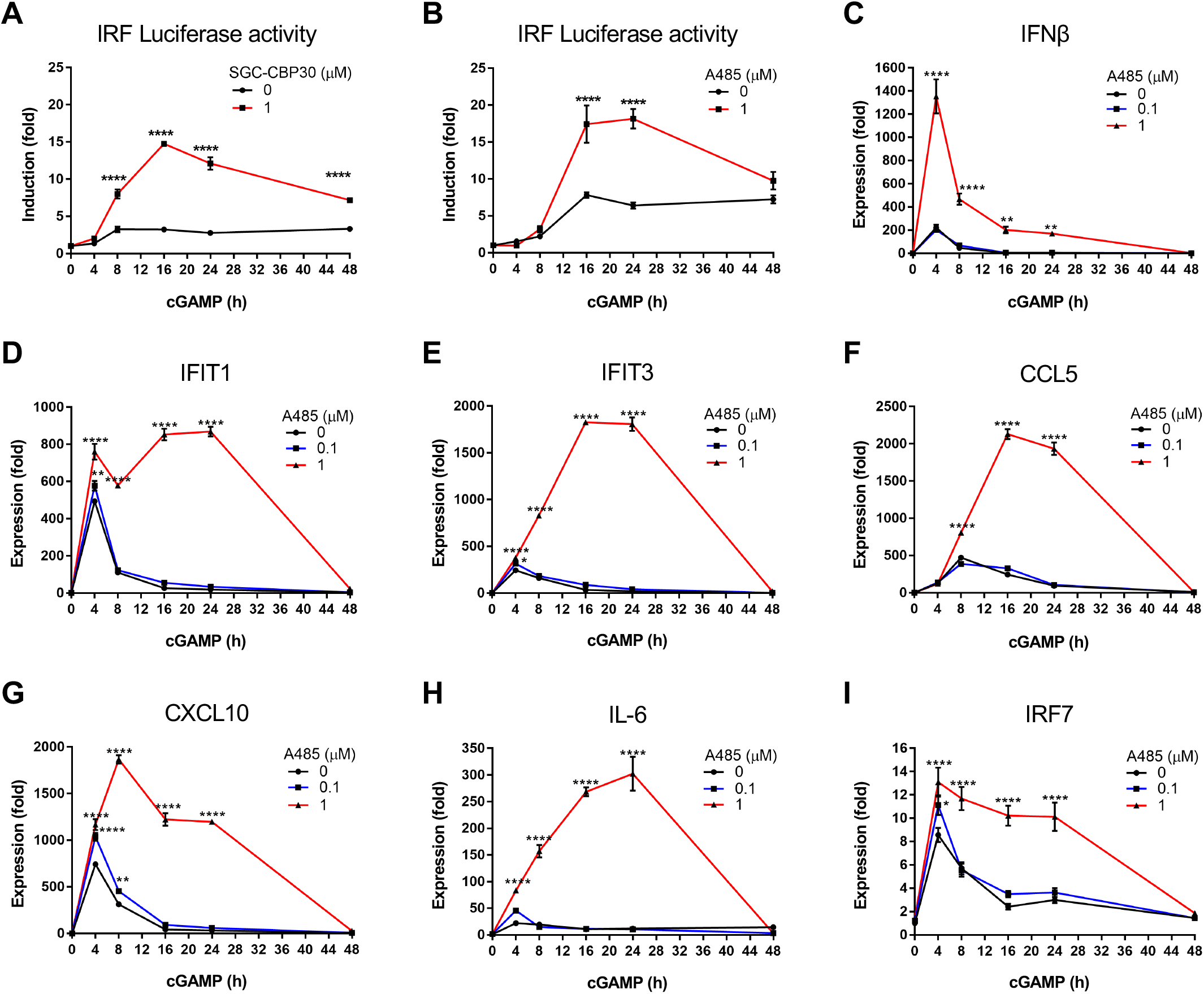
p300i enhances the transcription of innate immune genes. (**A-B**) IRF luciferase activity induced by cGAMP (2 µM) in Raw-Lucia ISG cells is enhanced by p300i SGC-CBP30 (1 µM) and A485 (1 µM), while p300i alone has no effect. (**C-I**) Pretreatment with A485 (0-1 µM) for 2h at each timepoint dose-dependently enhances and extends the transcription of innate immune responsive genes, including IFNβ, IFIT1, IFIT3 CCL5, CXCL10, IL-6 and IRF7 induced by cGAMP in Raw264.7 cells. A485 alone has no effect on the transcription of these genes measured by qRT-PCR. * *P*<0.05, ** *P*<0.01, **** *P*<0.0001.

Next, we measured the transcription of innate immune responsive genes at different timepoints of cGAMP stimulation, covering from 0 to 48h window. Treatment of macrophage-like Raw264.7 cells by p300i alone for either 2h or 50h did not induce the transcription of innate immune genes (shown at 0h timepoint). Pretreatment with A485 (1 µM), either 2h prior to cGAMP stimulation at each timepoint or simultaneously at the beginning of experiments, significantly boosted the cGAMP-induced transcription of innate immune genes, such as IFNβ, IFIT1, IFIT3, CCL5, CXCL10, and IL-6 in Raw264.7 cells, especially extending the active transcription period of these genes, so did SGC-CBP30 (Figure 1C-I and Figures S1, S2). To determine if the effect of p300i on STING-mediated innate immune responses was limited to macrophages, we tested non-immune cells, including 4T1 breast cancer cells and B16F10 melanoma cells. A485 (1 µM) also significantly enhanced the transcription of innate immune genes in response to cGAMP (5 µM) in these cells (Figures S3, S4). Thus, we discovered that p300i boosts the STING-mediated innate immune responses.

### p300i enhances IRF3 phosphorylation

To understand the impact of p300i on innate immune gene transcription, we first examined STING signaling in the cytoplasm, especially TBK1 phosphorylation (p-S172) and IRF3 phosphorylation (p-S396). TBK1 p-S172 correlates with its activated kinase activity (18). IRF3 phosphorylation at the C-terminal domain serine residues, such as p-S396 by TBK1, is a hallmark of IRF3 activation which allows its dimerization and subsequent nuclear translocation. At the baseline, TBK1 p-S172 and IRF3 p-S396 were largely undetectable by immunoblot. We first validated that cGAMP (2 µM) promptly induced p-S172 and IRF3 p-S396 in Raw264.7 cells but not in STING knockout (STING-KO) cells or pretreatment with TBK1 inhibitor GSK8612 (1 µM) (Figure 2A and Figure S5A-D). The cGAMP-induced TBK1 p-S172 peaked at 4h and subsequently declined but was still detectable at 48h in the cytosol. Comparatively, the cGAMP-induced IRF3 p-S396 was transient, peaked at 4h and then quickly disappeared in both cytosolic and nuclear fractions. Pretreatment with p300i alone or with a PROTAC degrader (dCBP1) alone did not induce any IRF3 phosphorylation (shown at 0h timepoint). However, with the presence of p300i and the degrader, TBK1 p-S172 was elevated from 4-48h of cGAMP stimulation and the duration of IRF3 p-S396 was extended to 48h timepoint in the nucleus (Figure 2B-C and Figure S5E). These results suggest that p300i promotes the cGAMP-induced phosphorylation and activation of TBK1 and IRF3.

**Figure 2.**
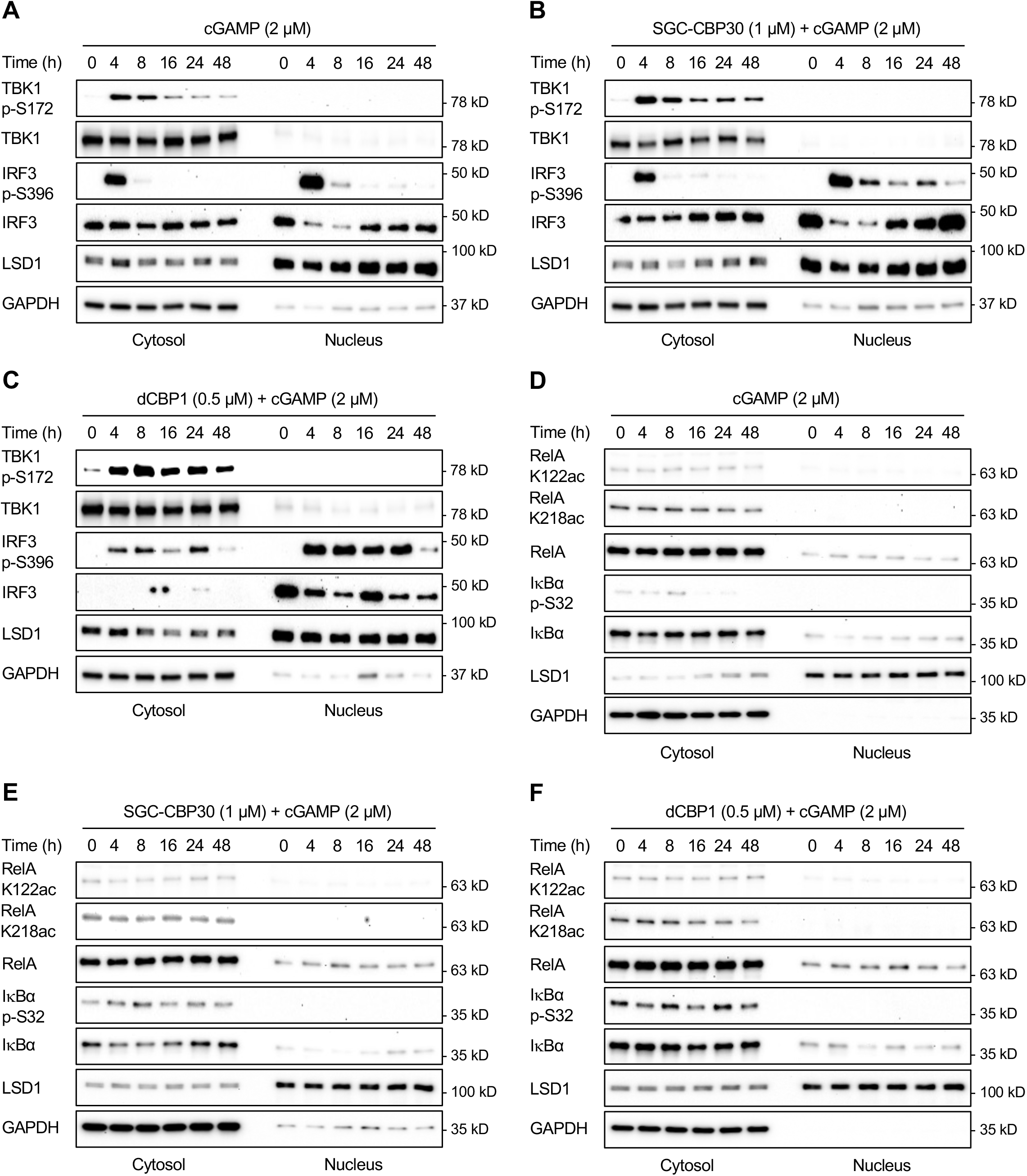
p300i promotes the phosphorylation of TBK1 and IRF3. (**A**) cGAMP (2 µM) induces TBK1 and IRF3 phosphorylation in Raw264.7 cells, shown by immunoblot. (**B**) p300i SGC-CBP30 (1 µM) enhances the cGAMP-induced phosphorylation of TBK1 and IRF3, while SGC-CBP30 alone (0h timepoint) has no obvious effects. (**C**) cGAMP induces IκBα phosphorylation and RelA nuclear translocation but no obvious changes in RelA acetylation in Raw264.7 cells, shown by immunoblot. (**D**) SGC-CBP30 (1 µM) enhances the cGAMP-induced IκBα phosphorylation and RelA nuclear translocation but does not affect RelA acetylation. LSD1: a nuclear marker; GAPDH: a cytosolic marker.

NF-κB proteins are normally sequestered in the cytoplasm by inhibitory partners such as IκB. In response to STING activation, TBK1 recruits and coordinates with IκB kinase to phosphorylate IκB proteins (e.g., IκBα) to permit IκB ubiquitination and degradation, which subsequently frees canonical NF-κB dimers (typically RelA/p50) to translocate to the nucleus (19, 20). The upregulated TBK1 activation by p300i could potentially facilitate NF-κB activation. We examined the impact of p300i on NF-κB activation, assessed mainly by the nuclear translocation of RelA. At the unstimulated state, nuclear RelA was low but detectable. cGAMP stimulation induced a mild nuclear translocation of RelA (Figure 2D). p300i and dCBP1 increased the presence of RelA in the nucleus after cGAMP stimulation (Figure 2E-F), which correlated with an enhanced IκBα phosphorylation. Furthermore, RelA acetylation at K122 and K218 were detected in the cytoplasm but not the nucleus. Neither cGAMP stimulation nor p300i obviously changed the levels of RelA K122ac and K128ac. No interactions between RelA and CBP or p300 was found by co-immunoprecipitation (co-IP) in Raw264.7 cells before or after cGAMP stimulation, indicating that CBP/p300 are not responsible for RelA acetylation and RelA acetylation unlikely plays a critical role in STING signaling either. Taken together, p300i mainly promotes the STING-mediated activation of IRF3.

### STING activation increases the presence of p300, not CBP, in the cytoplasm

The phosphorylation of IRF3 occurs in the cytoplasm after STING activation. Predominantly located in the nucleus (21), CBP/p300 presumably bind to IRF3 in the nucleus after IRF3 phosphorylation and nuclear translocation. This discrepancy cannot explain the robust promotion of IRF3 phosphorylation by p300i. We hypothesized that CBP and/or p300 might interact with TBK1 and/or IRF3 in the cytoplasm in response to STING activation. To test this hypothesis, we examined the subcellular localization of CBP or p300 by immunofluorescence in Raw264.7 cells, using antibodies that were designed to detect either CBP or p300 without cross reactions. In unstimulated cells, p300 predominantly resided in the nucleus. After cGAMP stimulation, the cytoplasmic p300 was significantly increased at 4h timepoint and subsequently reversed toward the baseline at 16h timepoint (Figure 3A-C). In contrast, there was only a trend of CBP increase in the cytoplasm at 4h timepoint of cGAMP stimulation, but not statistically significant (Figure 3D, E).

**Figure 3.**
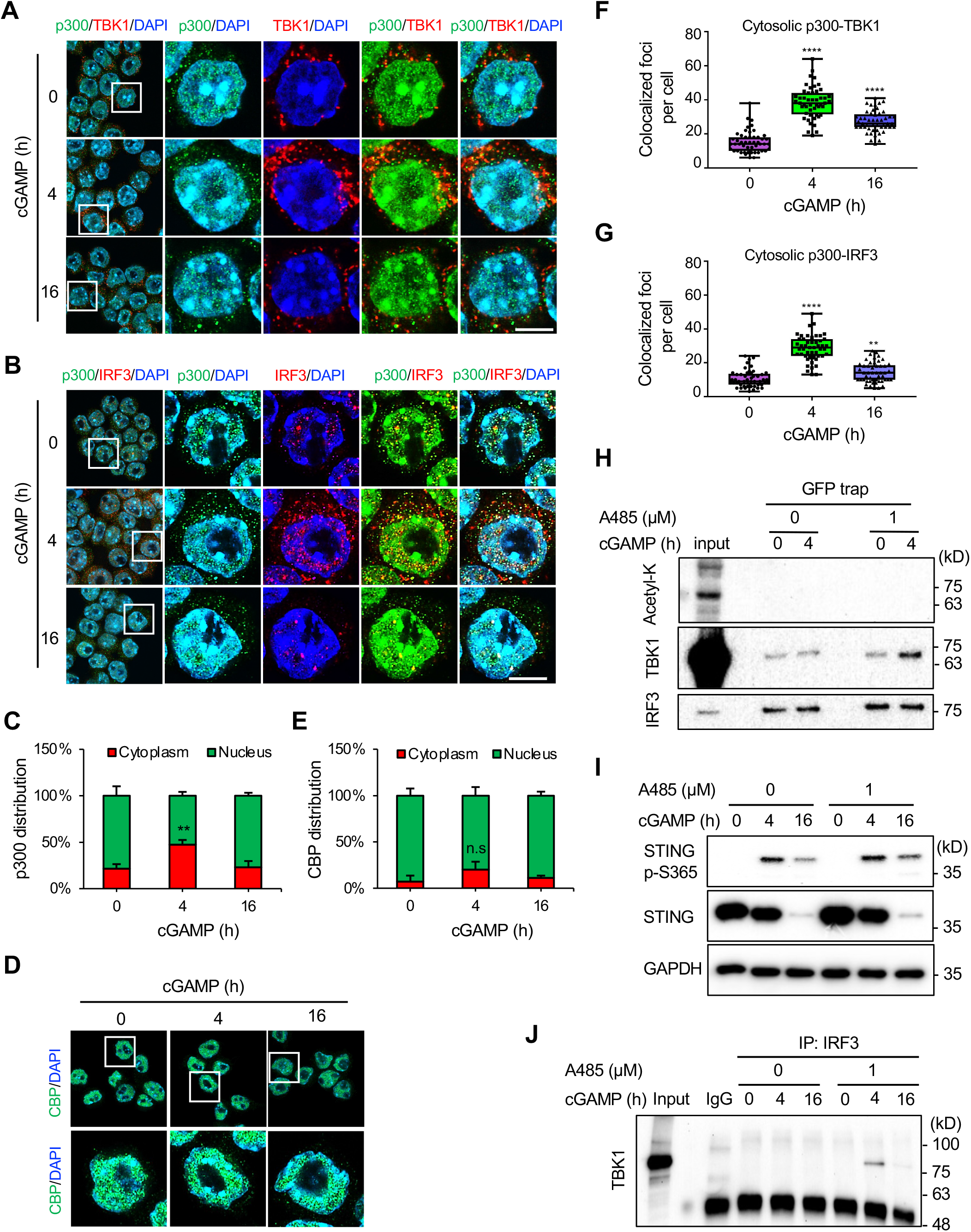
STING activation elevates cytoplasmic p300 to interrupt TBK1-IRF3 interactions. (**A-C**) cGAMP (2 µM) stimulation increases p300 in the cytoplasm of Raw264.7 cells, shown by immunofluorescence and image quantification. (**D-E**) cGAMP stimulation does not significantly change subcellular distribution of CBP. (**F-G**) Image quantification shows an increased colocalization of p300 with TBK1 or IRF3 in the cytoplasm at 4h timepoint of cGAMP stimulation, which subsequently reverses toward the bassline at 16h timepoint. (**H**) No acetylation of IRF3 is detected by GFP-Trap before or after cGAMP stimulation, with or without p300i A485 (1 µM) pretreatment. A485 enhances the interactions between TBK1 and GFP-IRF3. (**I**) A485 slightly increases STING phosphorylation (p-365) but has no obvious effects on STING degradation. (**J**) A485 enhances the interactions of endogenous TBK1 and IRF3, shown by co-IP. Scale bar=5 µm. n.s: not significant, ** p<0.01, *** p<0.001, **** p<0.0001.

The increased presence of p300 by STING activation indicates a possibility of its colocalization with TBK1 and/or IRF3 in the cytoplasm. In unstimulated cells, there was little colocalization of p300 with either TBK1 or IRF3 in the cytoplasm. At 4h timepoint of cGAMP stimulation, the colocalization of p300 with both TBK1 and IRF3 in the cytoplasm was prominent. Both colocalizations were subsequently decreased toward the baseline at 16h timepoint (Figure 3F, G). Taken together, these results suggest that STING activation increases the presence of p300, but not CBP, in the cytoplasm to colocalize with TBK1 and IRF3.

### p300i reinforces the interactions of TBK1 and IRF3

The physical proximity of p300 with TBK1 and IRF3 in the cytoplasm could potentially lead to TBK1 and/or IRF3 modifications. As an acetyltransferase, p300 is known to preferentially acetylate nuclear proteins including histones and non-histone proteins (22). Acetylation of cytoplasmic proteins by p300 is relatively rare but not excluded (23). We decided to detect potential acetylation of TBK1 or IRF3 by p300. Using co-IP, we found no evidence that TBK1 was acetylated in the cytosolic fractions of Raw264.7 cells before or after STING activation, witor without p300i A485 (Figure S5F). We then turned our attention to IRF3. To avoid IgG heavy chain interference with the immunoprecipitated IRF3 due to similar molecular weights, we overexpressed GFP-IRF3 by lentiviral gene delivery to Raw264.7 cells and enriched IRF3 from cytosolic fractions by GFP-Trap. Again, no IRF3 acetylation by p300 was detected (Figure 3H). Furthermore, the impact of p300i on STING was also minimal. A485 did not affect STING degradation and only slightly increased its phosphorylation (Figure 3I), which is likely a secondary effect of the upregulated TBK1 activation (7). These results indicate that neither TBK1 nor IRF3 is acetylated by p300 after STING activation.

To understand how p300i promotes IRF3 phosphorylation by TBK1, we examined interactions between TBK1 and IRF3 by co-IP. Without protein crosslinking, there were no visible co-IP bands before or after cGAMP stimulation. With A485 pretreatment, cGAMP stimulation caused a clearly visible co-IP band at 4h timepoint, which was diminished at 16h timepoint (Figure 3J). The strengthened interactions of TBK1 and IRF3 after STING action by p300i was subsequently validated by GFP-trap, although the exogenous GFP-IRF3 increased the level of baseline interactions (Figure 3H). These results suggest that p300i reinforces TBK1-IRF3 interactions after STING activation, resulting in higher IRF3 phosphorylation. Conversely, it is most likely that STING activation elevates cytoplasmic p300 to interrupt TBK1-IRF3 interactions and further inhibit IRF3 phosphorylation as a negative feedback regulation.

### Distinct roles of CBP and p300 in H3K27ac dynamics

To understand how p300i boosts the transcription of innate immune genes in response to STING activation, we assessed the acetylation of histone H3K27 which is one of lysine residues that are preferentially targeted by p300. Using immunoblot, we observed an unexpected H3K27ac reduction which was prominent at around 8h timepoint in Raw264.7 cells. H3K27ac was subsequently recovered toward the baseline at 48h timepoint (Figure 4A). cGAMP stimulation also caused a reduction in other histone acetylation marks, including pan H3ac, H4K16ac and pan H4ac (Figure 4A, B), suggesting global histone deacetylation. Rather than the expected histone acetylation by CBP/p300 after their presumed recruitment by the activated IRF3 to chromatin, STING activation appeared to lead to global histone deacetylation.

**Figure 4.**
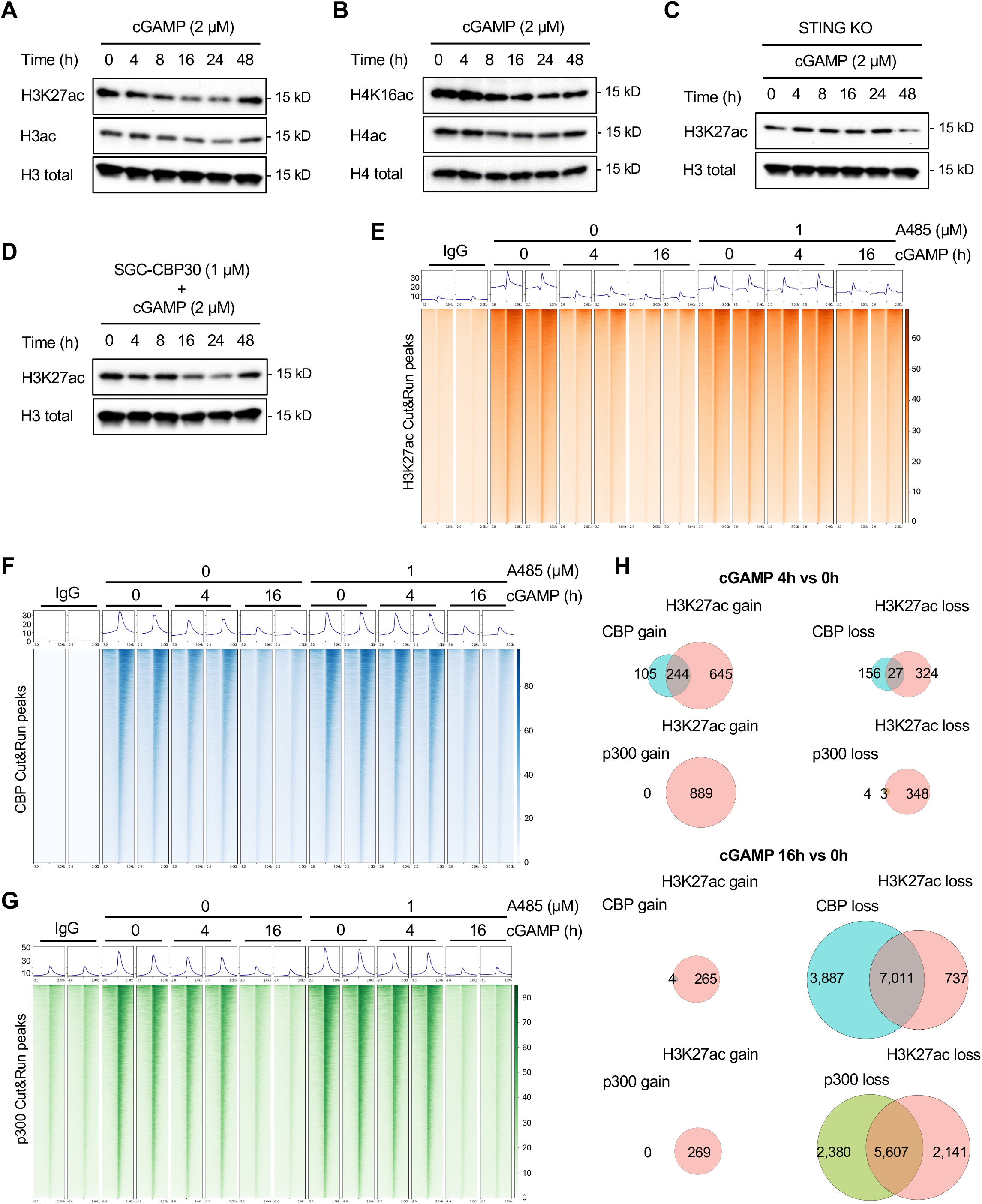
CBP and p300 play different roles in H3K27ac dynamics. (**A-B**) cGAMP (2 µM) decreases H3K27ac, H3ac, H416ac and H4ac in Raw264.7 cells which are subsequently reversed toward the baseline at 48h timepoint, shown by immunoblot. (**C**) In STING-KO Raw264.7 cells, cGAMP stimulation no longer decreases H3K27ac. (**D**) p300i SGC-CBP30 does not affect the cGAMP-induced H3K27ac reduction. (**E**) cGAMP (2 µM) stimulation diminishes H3K27ac peaks at 4 and 16h timepoints, shown by union peaks in Cut&Run heatmaps. p300i A485 (1 µM) does not change the trend of H3K27ac reduction after cGAMP stimulation but suppress H3K27ac across all timepoints. (**F-G**) cGAMP (2 µM) stimulation diminishes both CBP and p300 peaks at 16h timepoint, while having moderate impact at 4h timepoint, shown by union peaks in Cut&Run heatmaps. p300i A485 (1 µM) does not drastically change the pattern of CBP and p300 peaks induced by cGAMP stimulation. (**H**) At 4h timepoint, CBP, not p300, is associated with H3K27ac dynamics while at 16h timepoint, chromatin disassociation of CBP and p300 is linked to massive H3K27ac reduction, shown by correlation of statistically significant H3K27ac peaks with either CBP or p300 peaks.

In STING-KO Raw264.7 cells, cGAMP stimulation no longer reduced H3K27ac, indicating that H3K27ac reduction by cGAMP is dependent on STING (Figure 4C). Furthermore, p300i SGC-CBP30 (1 µM) pretreatment did not affect the cGAMP-induced H3K27ac reduction up to 24h timepoint nor H3K27ac recovery toward the baseline at 48h timepoint (Figure 4D). Overall, p300i pretreatment did not change the pattern of H3K27ac induced by STING activation. We then used Cut&Run to assess H3K27ac, CBP and p300 at different timepoints (0, 4, 16h) of STING activation in Raw264.7 cells. By examining union/all peaks, we verified the H3K27ac reduction after STING activation. With p300i, H3K27ac reduction after STING activation was delayed to 16h timepoint (Figure 4E). Comparatively, union peaks of CBP or p300 were slightly decreased at 4h timepoint of STING activation. However, both CBP and p300 peaks were largely diminished at 16h timepoint, indicating massive disassociation of CBP and p300 from chromatin (Figure 4F-G).

Further analysis of statistically significant peaks [p<0.05 after false discovery rate (FDR) correction] showed that, at 4h timepoint, there were 889 genes with upregulated H3K27ac peaks and 351 genes with downregulated H3K27ac peaks. At the same timepoint, there were 349 genes with upregulated CBP peaks and 183 genes with downregulated CBP peaks. Comparatively, there were no upregulated p300 peaks and merely 7 genes with downregulated p300 peaks. At 16h timepoint, there were almost no upregulated CBP or p300 peaks. In contrast, there were 10,898 genes with downregulated CBP peaks and 7,987 genes with downregulated p300 peaks, which were associated with massive H3K27ac peak reduction (n=7,748) (Figure 4H).

Combination of CBP, p300 and H3K27ac Cut&Run data indicated that, at 4h timepoint, the dynamic H3K27ac was correlated with CBP, not p300. In contrast, massive H3K27 deacetylation at 16h timepoint was correlated with the chromatin disassociation of CBP and p300 (Figure 4H). Together, these data suggest that CBP, not p300, underpins H3K27ac dynamics at early stages, while both CBP and p300 underlie H3K27 deacetylation at later stages, of STING activation.

### CBP, not p300, is co-recruited to chromatin with IRF3

To initiate the STING-mediated transcriptomic responses, the activated IRF3 is required to bind to chromatin. The genomic occupancy of IRF3 at different timepoints (0, 4, 16 h) of STING activation was assessed by Cut&Run. cGAMP (2 µM) stimulation induced a swift chromatin recruitment of IRF3 in Raw264.7 cells, which peaked at 4h timepoint and subsequently reversed toward the baseline at 16h timepoint, shown by union peak heatmaps (Figure 5A). Analysis of statistically significant peaks in comparison with the baseline showed that, at 4h timepoint, there were 502 genes with upregulated IRF3 peaks and 93 genes with downregulated IRF3 peaks. At 16h timepoint, there were no genes with upregulated IRF3 peaks and 128 genes with downregulated IRF3 peaks. Overall, STING activation causes a relatively short-term chromatin engagement of IRF3.

**Figure 5.**
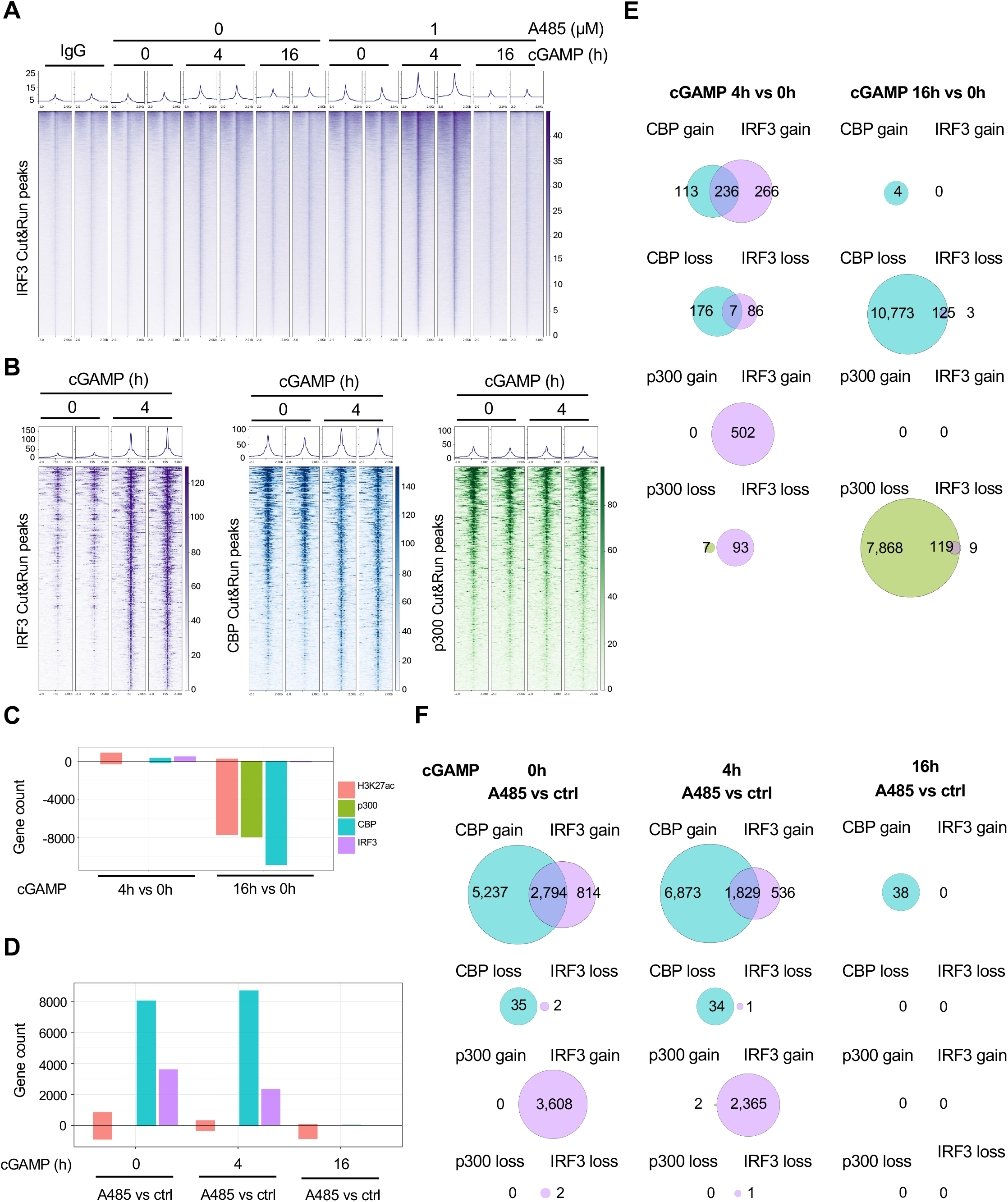
Chromatin co-recruitment of IRF3 and CBP, but not p300. (**A**) cGAMP (2 µM) induces IRF3 chromatin recruitment in Raw264.7 cells at 4h timepoint, which is subsequently reversed toward the baseline at 16h timepoint, shown by union peaks in Cut&Run heatmaps. p300i A485 (1 µM) markedly enhances IRF3 chromatin recruitment at 4h timepoint. (**B**) Correlation of IRF3 and CBP chromatin recruitment at 4h timepoint when no obvious p300 chromatin recruitment is found, shown by statistically significant Cut&Run peaks. (**C**) Bar chart shows numbers of gene that are associated with statistically significant H3K27ac, p300, CBP, IRF3 peaks in timewise comparisons. (**D**) Pairwise comparisons of statistically significant Cut&Run H3K27ac, p300, CBP, IRF3 peaks that are affected by A485. (**E**) At 4h timepoint, CBP, not p300, is associated with IRF3 dynamic chromatin engagement while at 16h timepoint, chromatin disassociation of IRF3 is mainly linked to CBP, not p300, shown by correlation of statistically significant H3K27ac peaks with either CBP or p300 peaks. (**F**) A485 increases IRF3 peaks at 0 and 4h, but not 16h of cGAMP stimulation. Pairwise correlations of statistically significant peaks show that A485 enhances the chromatin recruitment of IRF3 and CBP, but not p300, at 0h and 4h timepoints, while having minimal effect at 16h timepoint. ctrl: control.

At 4h timepoint, IRF3 and CBP, but not p300, were found to recruit to chromatin, shown by heatmaps of statistically significant peaks (Figure 5B). Combination of IRF3 with CBP or p300 data showed a correlation between IRF3 and CBP, but not p300. At 4h timepoint, there were 236 genes with upregulated IRF3/CBP overlapping peaks and 7 genes with downregulated IRF3/CBP overlapping peaks. However, no genes were found to contain either upregulated or downregulated IRF3/p300 overlapping peaks (Figure 5C, E). CBP, not p300, appeared to co-recruit to chromatin with IRF3 after STING activation. As described above, CBP, not p300, was associated with H3K27ac upregulation at 4h timepoint of cGAMP stimulation. Together, these data suggest that CBP, not p300, acts as a transcription coactivator for IRF3 in cGAS-STING pathway.

At 16h timepoint, there were 125 genes with downregulated IRF3/CBP overlapping peaks and 119 genes with downregulated IRF3/p300 overlapping peaks, while no genes were found to contain upregulated IRF3/CBP overlapping peaks or IRF3/p300 overlapping peaks (Figure 5C, E), indicating that the disengagement of IRF3 from chromatin was correlated with the chromatin disassociation of CBP and p300, which was also linked to massive H3K27 deacetylation as shown above. Taken together, these data suggest that, at later stages of STING activation, nearly all CBP and p300 were detached from chromatin, which is associated with global histone deacetylation and the chromatin eviction of IRF3.

### p300i boosts IFN responses by promoting IRF3 chromatin recruitment

Without cGAMP stimulation, p300i A485 alone increased the binding of IRF3 and CBP to the chromatin, shown as 2,794 genes with upregulated IRF3/CBP overlapping peaks. In contrast, A485 had no obvious impact on the chromatin association of p300 before or after STING activation. At 4h timepoint, A485 and cGAMP combination further enhanced the chromatin recruitment of IRF3 and CBP, shown as 1,829 genes with upregulated IRF3/CBP overlapping peaks, compared to cGAMP alone group at the same timepoint. At 16h timepoint, there were no differences in IRF3 chromatin engagement between cGAMP alone group and the combination group (Figure 5D, F). These data suggest that p300i promotes the chromatin recruitment of IRF3 and CBP, but not p300, which also correlates with the delayed H3K27ac reduction, at early stages of STING activation.

Using bulk RNA-seq, we validated that cGAMP induced a rapid shift in the transcriptome. Compared to the baseline (0h), there were 594 transcripts upregulated and 94 transcripts downregulated (log_2_-fold change threshold at ±1) at 4h timepoint. At 16h timepoint, there were 621 transcripts upregulated and 128 transcripts downregulated (Figure 6A). Pairwise comparisons at each timepoint showed that A485 alone largely suppressed the transcriptome, shown as 212 downregulated transcripts and 18 upregulated transcripts at 0h timepoint. At 4h timepoint of cGAMP stimulation, A485 downregulated 231 transcripts (no signature innate immune genes found) and upregulated 41 transcripts, including IFNβ. At 16h timepoint, A485 downregulated 977 transcripts and upregulated 1,003 transcripts, including IFNβ and ISGs (Figure 6B). Pathway analysis showed that A485 suppressed inflammatory responses, while promoting IFN responses at early hours of cGAMP stimulation. At 16h timepoint, A485 dramatically boosted IFN responses, moderately enhanced inflammatory responses and markedly suppressed cell cycle pathways (Figure 6C).

**Figure 6.**
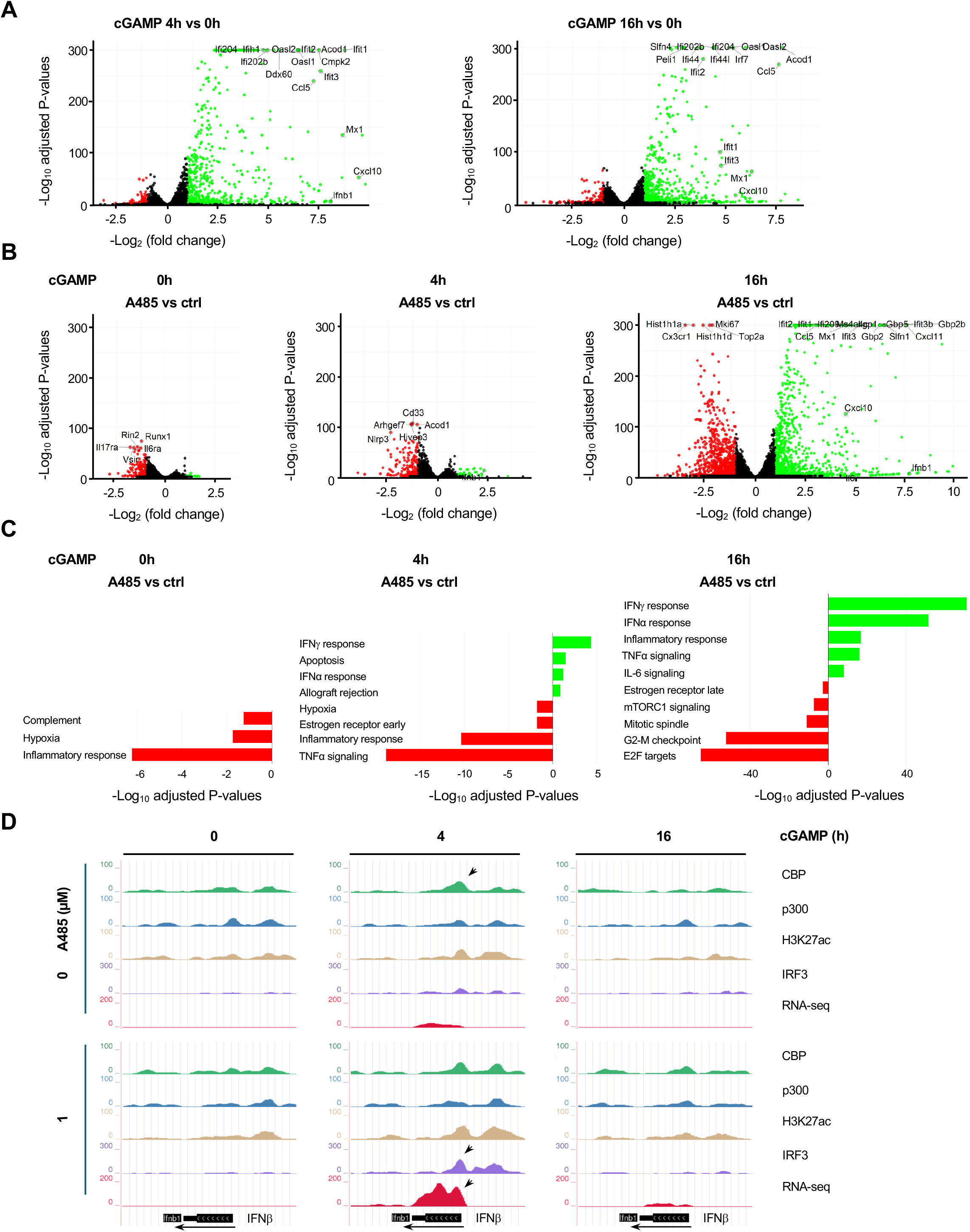
p300i enhances IFN response by promoting IRF3 chromatin engagement. (**A**) cGAMP (2 µM) induces robust transcriptomic responses at 4h and 16h timepoints, shown by RNA-seq volcano plots. (**B**) p300i A485 (1 µM) suppresses the transcriptome at the baseline. A485 largely suppresses the cGAMP-modified transcriptome while increasing IFNβ transcription at 4h timepoint. At 16h timepoint, A485 dramatically promotes the cGAMP-induced transcription of IFNβ and ISGs. (**C**) Pathway analysis shows that A485 enhances the cGAMP-induced IFN responses. (**D**) Sequencing traces show that cGAMP induces the transcription of IFNβ at 4h timepoint, which correlates with the upregulated peaks of IRF3, H3K27ac and CBP (arrow) at the promoter region where no obvious changes in p300 peaks are found. A485 increases the transcription of IFNβ at 4h timepoint (arrow) and extends IFNβ transcription at 16h timepoint, which correlates with the markedly elevated IRF3 peaks at the promoter region (arrow).

We then examined sequencing traces of individual genes, especially the signature genes of STING-mediated innate immune responses. There were no RNA reads aligned to IFNβ at the baseline. At 4h timepoint of cGAMP stimulation, an RNA-seq peak emerged which subsequently disappeared at 16h timepoint. Correspondingly, cGAMP induced IRF3 peaks, CBP peaks and H3K27ac peaks, but not p300 peaks in the promoter region of IFNβ at 4h timepoint, of which all vanished at 16h timepoint (Figure 6D). The transcription of IFNβ was correlated with chromatin engagement of IRF3 and CBP, but not p300, in its promoter.

p300i A485 alone did not induce IFNβ transcription. With A485 pretreatment, RNA-seq peaks of IFNβ were much higher at 4h timepoint of cGAMP stimulation, which were decreased but still visible at 16h timepoint. At 4h timepoint, A485 markedly increased IRF3 peaks, while having no obvious effect on CBP, p300 and H3K27ac peaks (Figure 6D). The enhanced transcription and IRF3 chromatin engagement by A485 were also found in other innate immune responsive genes, such as CXCL10 (Figure S6). Taken together, these analyses indicate that p300i boosts innate immune responses, especially IFN responses, to STING activation by promoting IRF3 chromatin recruitment.

## DISCUSSION

cGAS-STING pathway, similar to other signal transduction pathways, consists of three major segments, i.e., reception, transduction, and response. Intense research of more than a decade from many groups has yielded a much clearer picture about how dsDNA fragments activate the receptor cGAS to synthesize the second messenger cGAMP (i.e., reception) and how the adaptor STING transduces cGAMP to IRF3 phosphorylation and activation (i.e., transduction), as well as the regulation of these processes (24, 25). However, transcriptomic responses to STING activation, which are driven primarily by IRF3, remain not fully understood.

Overactivation of cGAS-STING pathway, especially to self-DNA, can lead to autoimmune and autoinflammatory disorders, such as those caused by impaired digestion of chromatin (PMID: 23666765, PMID: 27293190). Many layers of negative regulation have evolved to restrain STING signaling, such as cytosolic DNA degradation, cGAS nuclear sequestration and STING degradation, to prevent cGAS-STING overactivation (26, 27). Results of this study suggest a new layer of negative regulation of STING signaling by p300, and most likely not CBP. In unstimulated cells, CBP and p300 are predominately located in the nucleus. STING activation increases the presence of p300, not CBP, in the cytoplasm, which is likely caused by the diminished AMPK-catalyzed phosphorylation of p300 (p-S89), an essential modification for p300 nuclear translocation (28–30). Unfortunately, lack of reliable antibodies, due to commercial product discontinuation, prevents us from testing the potential mechanism underpinning cytoplasmic retention of p300 by STING activation in the current study. Nevertheless, the elevated cytoplasmic p300 appears to interrupt interactions between TBK1 and IRF3 to block IRF3 phosphorylation. It is noteworthy that cytoplasmic acetyltransferases KAT2a/b have been found to also inhibit TBK1 activation in cells after RNA viral infections (31), indicating a potential shared mechanism. On the other hand, p300i markedly enhances TBK1-IRF3 interactions, promotes IRF3 phosphorylation, and boosts the transcription of innate immune responsive genes. Cytoplasmic p300 thus is a previously unknown suppressor of IRF3 activation in cGAS-STING pathway.

CBP and p300 have been thought to be redundant in cGAS-STING pathway. Both are potentially recruited by IRF3 as transcription coactivators. However, studies show that CBP and p300 are not always redundant and can wield non-overlapping functions (32–35). Our experiments indicate that only CBP, not p300, is co-recruited to chromatin with IRF3 after STING activation. Furthermore, at early stages of STING activation, H3K27ac dynamics, especially the upregulated H3K27ac, are associated with the chromatin-bound CBP, but not p300. For the first time, we have identified that CBP, not p300, acts as a transcription coactivator for IRF3 in cGAS-STING pathway.

STING-mediated innate immune responses are often transient, which can be partially explained by the negative regulations of STING signaling in the cytoplasm. One key issue underpinning the transient innate immune responses is to explain how IRF3, and potentially other transcription factors, are evicted from chromatin to terminate their transcription programs. Data analyses of current research indicate that, at later stages of STING activation, nearly all CBP and p300, including the ones that are chromatin-bound prior to cGAMP stimulation, are detached from chromatin. The massive detachment of CBP and p300 from chromatin causes global histone deacetylation that is associated with the chromatin eviction of IRF3, which can consequently terminate its transcription programs.

Using p300i as a tool to wedge into cGAS-STING pathway, our study reveals distinct roles of CBP and p300 in cGAS-STING pathway. In the cytoplasm, p300, not CBP, is a suppressor of STING signaling. In the nucleus, CBP, not p300, serves IRF3 as a transcription coactivator for the transcription of IFNs and ISGs. At later stages of STING activation, chromatin disassociation of both CBP and p300 causes global histone deacetylation, likely leading to the chromatin eviction of IRF3 and the termination of its transcription programs. Furthermore, p300i boosts STING-mediated innate immune responses primarily by promoting IRF3 activation and enhancing IRF3 chromatin recruitment, while inhibiting CBP in the nucleus, suggesting that the inhibitory role of p300 on IRF3 by limiting its activation outweighs the transcription coactivator role of CBP.

More work is required to further dissect mechanisms underpinning the roles of CBP and p300 in cGAS-STING pathway. For example, it remains elusive how cytoplasmic p300 inhibits TBK1-IRF3 interactions to prevent IRF3 phosphorylation. Our data show that the inhibition is dependent on p300 functions since p300i strongly enhances TBK1-IRF3 interactions. However, no acetylation of TBK1 or IRF3 by p300 has been identified, suggesting potential involvement of an unknown substrate of p300 in the regulation of TBK1-IRF3 interactions. Moreover, factors that trigger massive disassociation of CBP and p300 from chromatin at later stages of STING activation are unknown. These and many other unresolved questions surrounding the distinct roles of CBP and p300 in cGAS-STING pathway are warranted for further investigations.

Intriguingly, gain-of-function mutations in p300 cause Menke-Hennekam syndrome which has a common feature of recurrent infections, particularly in the upper respiratory tract (36). Whereas loss-of-function mutations in CBP lead to Rubinstein-Taybi syndrome which frequently shows immunological deficiency and an increased rate of respiratory infection (37, 38). How gain-of-function mutations in p300 and loss-of-function mutations in CBP lead to similar immunological deficiency have puzzled the field for some time. These syndromes have not yet been linked to cGAS-STING pathway. Our discovery of distinct roles of CBP and p300 in cGAS-STING-mediated innate immunity opens a path to elucidate pathogenic mechanisms of these disorders. It is plausible that the gain-of-function mutations in p300 strengthens its suppressor role on STING signaling, while the loss-of-function mutations in CBP disrupts its transcription coactivating function, both of which inhibit STING-mediated innate immunity, leading to immunological deficiency.

In summary, our findings suggest that p300, not CBP, is a suppressor of STING signaling in the cytoplasm. In contrast, CBP, not p300, acts as a transcription coactivator in the nucleus. Massive chromatin disassociation of CBP and p300 at later stages of STING activation causes global histone deacetylation, likely leading to the chromatin eviction of IRF3 and the termination of its transcription programs. p300i boosts STING-mediated innate immune responses, which can be used as a potential therapeutic strategy to treat infectious diseases and cancer.

## METHODS

### Sex as a biological variable

This study used commercially available cells derived from both male and female patients in the experiments. Sex is not a biological variable.

### Cell culture

Raw264.7 cells and Raw-Lucia ISG cells were purchased from ATCC and InvivoGen respectively, between the year of 2019 to 2021 without further authentication. B16F10 cells and 4T1 cells were also purchased from ATCC. Frozen cells were newly thawed from low (3–10) passages. Mycoplasma was tested using PlasmoTest mycoplasma detection kits (ThermoFisher Scientific, #4460626) and only negative cells were included in all experiments. All cells were maintained under a 5% CO_2_ atmosphere in Dulbecco’s Modified Eagle’s Medium (ThermoFisher Scientific, #11995065) supplemented with 10% heat-inactivated FBS, 100 Units/ml penicillin, and 100 µg/ml of streptomycin.

### Luciferase assay

RAW-Lucia ISG cells were derived from RAW 264.7 cell line by stable integration of an IRF-inducible Lucia luciferase reporter construct, which is under the control of an ISG54 promoter in conjunction with five ISRE. Raw-Lucia ISG cells were seeded in 96-well plates (1×10^4^ cells/well) for 24h and pretreated with p300i SGC-CBP30 (1 µM) (Selleck Chemicals, #S7256) or A485 (1 µM) (Selleck Chemicals, #S8740) for 1 h. After treatment with 2 µM cGAMP (InvivoGen, #tlrl-nacga2srs-05) for 0-48h, cell lysates were harvested for the evaluation of IRF induction using QUANTI-Luc reagent (InvivoGen, #rep-qlc4lg1) and a Bio-Tek Synergy2 Plate Reader. Relative IRF induction activity was calculated as luminescence fold changes over the unstimulated controls.

### Quantitative RT-PCR

Raw264.7 cells, B16F10 cells and 4T1 cells were treated with SGC-CBP30 (0-1 µM) or A485 (0-1 µM) for either 2h prior to cGAMP stimulation at each timepoint or simultaneously at the beginning of experiments for 50h. Total RNA was extracted from cells using QIAzol reagent (Qiagen, #79306). cDNA was synthesized from 1 µg of total RNA with SuperScript III First-Strand Synthesis System (ThermoFisher Scientific, #18080051). Real-time quantitative PCR was performed with 2 µL of 1:10 diluted cDNA per reaction on a Quant studio 12K Flex real-time PCR system (ThermoFisher Scientific) using PowerUp Sybr Green Master Mix (ThermoFisher Scientific, #A25742). Primers were designed to span introns (Table S1). Relative gene expression was determined using the 2^−ΔΔ*C*T^ method.

### STING knockout

To generate STING-KO RAW 264.7 cells, custom sgRNA were designed (5′-C*A*C*CUAGCCUCGCACGAACU-3′) and procured using Alt-R-CRISPR/Cas9 platform (IDT). Cas9-gRNA ribonucleoprotein complexes were assembled using 140 pmol sgRNA and 40 pmol EnGen® Spy Cas9 HF1 (New England Biolabs, #M0667M) and incubated for 10 min at room temperature. Subsequently, 8×10^5^ cells were suspended in the RNP complex in 80 µL of nucleofector solution. Nucleofection was performed using SF cell line-Nucleofector kit (Lonza, #V4XC-2012) and preprogrammed pulse protocol DS-136 in 4D-Nucleofector system (Lonza). After recovery, cells were plated in 96-well plates at an average of 0.5 cells per well for generating monoclonal lines using Poisson distribution. Sanger sequencing was used to validate STING KO and to exclude potential off target editing by examining top five possible off-target *loci* provided by the CRISPR design tool. The sequence-verified clones were expanded for several passages to confirm cell integrity. STING KO was further validated at mRNA and protein levels by qRT-PCR and immunoblot.

### Immunoblot

Cells were stimulated with 2 µM cGAMP for 0, 4, 8, 16, 24 and 48h. After treatment, cells were rinsed twice with ice cold PBS and collected by centrifuging at 600× g for 5 min. Cell pellets were dislodged with 1 mL nuclear extraction buffer (EpiCypher, #21-1026a) containing protease inhibitors and spermidine for 10 min on ice. Nuclei were precipitated by centrifugation at 600× g for 3 min and supernatants were collected as cytosolic fractions. Nuclear pellets were dissolved in 0.5 mL IP lysis buffer (ThermoFisher Scientific, #87788) containing protease inhibitors at 4 °C to incubate for 30 min on a rotator, which was followed by centrifugation at 14,000× g for 10 min to separate supernatants as nuclear fractions. Both cytosolic and nuclear fractions were quantified using BCA protein assay kit (ThermoFisher Scientific, #55864). Samples were subjected to SDS-PAGE and transferred to nitrocellulose membranes using a Turbo blot transfer unit (Bio-Rad). Transfer efficiency was determined by Ponceau S staining (ThermoFisher Scientific, #A40000279). Membranes were incubated with blocking solution (TBS containing 0.1% Tween 20 and 5% nonfat dry milk) and were probed with different primary antibodies, including anti-H3K27ac (Active Motif, #39685, 1:3,000), anti-H4K16ac (Active Motif, #13534, 1:3,000), anti-H3ac (Active Motif, #39139, 1:3,000), anti-H4ac (Active Motif, #39925, 1:3,000), anti-TBK1 (Cell Signaling Technology, #3013, 1:1,000), anti-TBK1 p-S172 (Cell Signaling Technology, #5483, 1:1,000), anti-IRF3 (Cell Signaling Technology, #4302, 1:1,000), anti-IRF3 p-S396 (Cell

Signaling Technology, #4947, 1:1,000), anti-RelA (Cell Signaling Technology, #8242, 1:1,000), anti-RelA K122ac (Signalway Antibody, #HW148, 1:1000), anti-RelA K218ac (Signalway Antibody, #HW122, 1:1000), anti-IκBα (Cell Signaling Technology, #4814, 1:1,000), anti-IκBα p-S32 (Cell Signaling Technology, #2859, 1:1,000), anti-LSD1 (Cell Signaling Technology, #4064, 1:1,000), anti-GAPDH (Cell Signaling Technology, #97166, 1:1,000), for overnight at 4°C. After washing, membranes were incubated with HRP-conjugated secondary antibodies (Cell Signaling Technology, anti-mouse IgG, #7076, 1:10,000; anti-rabbit IgG, #7074, 1:10,000) for 1h at room temperature. Protein bands were visualized by SuperSignal West Pico PLUS Chemiluminescent Substrate (ThermoFisher Scientific, #34580) and captured by a FluorChem E chemidoc machine (Bio-Techne).

### Immunofluorescence

Cells were seeded on chamber slides and stimulated with 2 µM cGAMP for 0, 4 and 16h. Cells were then fixed with 4% paraformaldehyde for 10 min at room temperature and permeabilized for 5 min with 0.2% Triton X-100. After blocking with 5% BSA for 1h, slides were incubated with primary antibodies: anti-p300 (Cell Signaling Technology, #54062, 1:200); anti-CBP (Cell Signaling Technology, #7389, 1:200); anti-p300 (Cell Signaling Technology, #54062, 1:200) and anti-TBK1 (Cell Signaling Technology, #51872, 1:300) at 4°C overnight and followed by respective secondary antibody, including Alexa fluor 488 donkey anti-rabbit IgG (Life Technologies, #A11008, 1:400) and Alexa fluor 568 goat anti-mouse IgG (Life Technologies, #A11004, 1:400) for 2h. Each step was preceded by a three-time wash in PBST (0.2% Triton X-100 in PBS). Slides were mounted with DAPI containing antifade diamond mounting media. Images were captured by a Zeiss LSM980 confocal microscope with super resolution mode under 63× oil objective with 3× zoom for clear colocalization signals between two fluorochromes. Intensity of CBP/p300 signal in cytoplasm and nucleus was quantified for multiple areas using ImageJ selection tools at different timepoints (n=3 slides/timepoint). The colocalized spots of CBP/p300 with TBK1 or IRF3 were manually counted per cell for 50 cells for each slide (n=3 slides/timepoint).

### Co-immunoprecipitation

Cells were treated with 2 µM cGAMP for 0, 4 and 16 h. After treatment, cytosolic and nuclear fractions were extracted as described in Immunoblot. 1 mg of cytosolic proteins were incubated with 1-2 µg IRF3 antibody (Cell Signaling Technology, #4302) or the same amount of IgG overnight by rotating at 14 rpm at 4°C. Captured immunocomplexes in 1× laemmli buffer were boiled at 95°C for 5 min, electrophoresed in SDS-PAGE gels, and subjected to immunoblot. Alternatively, samples were incubated with IRF3 antibody or IgG crosslinked to Protein A/G magnetic beads using antibody crosslinking kit (ThermoFisher Scientific, #88805) overnight at 4°C followed by washes and elution. Eluted samples were then denatured and separated by SDS-PAGE and transferred to nitrocellulose membrane for immunoblot analysis using primary antibodies, including p300 (Cell Signaling Technology, #54062, 1:1,000), CBP (Cell Signaling Technology, #7389, 1:1,000), acetyl-K (Invitrogen, #MA1-2021, 1:1,000), anti-TBK1 (Cell Signaling Technology, #3013, 1:1,000) and IRF3 antibody (Cell Signaling Technology, #10949, 1:1,000).

### GFP-trap

RAW 264.7 cells were stably transduced with lentivirus hosting GFP tagged-IRF3 from Origene (#MR206641L4V). Positive cells were selected by puromycin resistance and single cell clones were established and maintained under puromycin. When reached to around 60% confluency in 150 mm dishes, cells were treated with cGAMP at 2 µM (InvivoGen) for 0, and 4h. Post-stimulation cells were carefully rinsed twice with ice cold PBS and collected by centrifuging at 600g for 5 min. Cell pellets were dislodged with 1 ml nuclear extraction buffer (Epicyhper #21-1026a) containing protease inhibitors and spermidine for 10 min on ice followed by centrifugation at 600g for 3 min to pellet nuclei. The supernatant was carefully collected into a new tube labelled cytosolic fraction without touching the pellet. Cytosolic lysate was quantified for protein using BCA protein assay kit. 1 mg of protein was incubated with 25 µl GFP nanobody coupled magnetic particles M-270 from Protein Tech (#gtd-20) overnight at 4°C. After rigorous washes, the trapped proteins were eluted in 1× laemmli sample buffer supplemented with BME by heating at 95°C for 5 min. Eluted samples were separated by SDS-PAGE and subjected to immunoblot using anti-TBK1, anti-Acetyl-K and anti-IRF3 antibodies.

### Cut&Run

Cut&Run was performed using 500,000 cells (n=2/group) stimulated with 2 µM cGAMP for 0, 4 and 16h. Cells were immobilized on concanavalin A–coated magnetic beads and nuclei were extracted using pre-nuclear extraction buffer (EpiCypher, #21-1026a). The extracted nuclei were processed with a CUT&RUN kit (EpiCypher, #14-1048) according to the manufacturer’s instruction. After permeabilization, the resuspended nuclei were incubated with antibodies against p300 (Cell Signaling Technology, #54062, 1:50), CBP (Cell Signaling Technology, #7425, 1:50), H3K27ac (EpiCypher, #13-0059, 1:50), IRF3 (Cell Signaling Technology, #4302, 1:50), or IgG (EpiCypher, #13-0042, 1:50) as negative controls. The enriched DNA was quantified using a Qubit fluorometer and processed to generate sequencing libraries using CUT&RUN Library Pre Kits (EpiCypher, #14-1001). Libraries were then pooled and analyzed using tape station for the fragment size followed by sequencing on an Illumina NovaSeq X sequencer to generate 10 million unique reads per sample.

Universal adapter sequences were trimmed from all reads using trim_galore. The trimmed reads were then aligned to the mouse genome mm10 using bwa (v0.7.17). Peak calling was performed using MACS2 (v2.2.7.1) with default parameters. The final peak list was composed of the sample peak list subtracting the set of peaks called in the IgG samples. Differential peaks were calculated using narrowPeak files produced by MACS2 and the aligned bam files by the R package DiffBind (v3.4.11). DESeq2 was used as the internal statistical model from within DiffBind. To create the heatmap, bigwig files were created using the internal spike in control to normalize the coverage using deeptools (v3.5.3) bamCoverage function. These bigwig files were then used as input to the computeMatrix function using the peaks that were found by MACS2. This produced a matrix file that was then fed into plotHeatmap (also within deeptools) where the median was used as the summary plot type. The final heatmaps generated show a union of all peaks found by MACS2, and other heatmaps consist of statistically significant (FDR<0.05) peaks. Gene lists were constructed using the R package ChIPSeeker (v.1.42.1) to annotate the genomes using mm10 as a reference.

### RNA-seq

Total RNA was extracted from Raw264.7 cells, which had been treated with 2 µM cGAMP for 0, 4 and 16h (n=2/group), using QIAzol reagent. A Bioanalyzer 2000 was used to measure the quality of RNA. All samples’ RNA integrity numbers (RIN) were above 9. After ribosomal RNA (rRNA) was depleted, sequencing libraries were ligated with standard Illumina adaptors and subsequently processed by a NovaSeq X sequencer. Universal adapter sequences were trimmed from raw fastq files using trim_galore. Alignment was performed against the mm10 genome using the STAR aligner (v.2.5.2a) with default parameters. The quantification of gene expression was performed by STAR as well using the Ensembl gtf file (v91). Gene expression count files were rearranged into matrices to determine differential expression by DESeq2. The differentially expressed genes (DEGs) were then filtered by a log_2_FC of ±1. Subsequently, volcano plots were created with R. The filtered DEGs were also fed into Enrichr (https://maayanlab.cloud/Enrichr/) for enrichment analysis (39–41).

### Integration of Cut&Run and RNA-seq

IRF3 and NF-κB identified in Cut&Run analysis were fed into ChIPseeker (v1.42.1) for annotation (42). All promoter annotations were merged to form the “Promoter”, all exon/intron 5’UTR/3’UTR were merged to form the “Gene body” annotation, and the rest were considered “Intergenic”. These final list of annotated genes were compared and overlapped with DEGs from RNA-seq data. Bigwig files that were created during Cut&Run analysis were used to create custom UCSC genome browser tracks in a local hub.

### Statistical analysis

Statistical analyses were performed using Microsoft Excel and GraphPad Prism Software. Data was analyzed by unpaired two-tailed Student’s *t-*test for two-group comparisons or by analysis of variance (*ANOVA*) test for more than two groups. Results with *P<0.05* were considered statistically significant. For Cut&Run and RNA-seq analysis, CBP, p300, or IRF3 peaks and DEGs with adjusted *P<0.05* were considered statistically significant.

## Supplemental Information

This article contains supplementary information comprised of supplementary figures and a table.

## Data Availability Statement

RNA-seq and Cut&Run sequencing data were deposited in the NCBI’s GEO database (RNA-seq: GSE344545; Cut&Run: GSE344665). Other data generated or analyzed during this study are included in this article and its supplementary information files.

## Acknowledgments

We thank Dr. Jose Lutzky and Dr. Susan Kesmodel of University of Miami for providing additional funds to support this work. We also thank Dr. Felipe Beckedorff for technical support and Ms. Erna Stoddart for administrative assistance.

## Author Contributions

Conceptualization: G.W.; Experimental design and supervising: G.W. and M.T.; Data collection: K.G. and M.F.Z.; Data analysis: D.V.B., K.G., G.W. and M.T.; Funding acquisition: G.W.; Manuscript drafting: G.W., K.G., and M.T., Manuscript review and editing: K.G., D.V.B., M.F.Z, M.T. and G.W.

