## Supplemental Figures and Table 1. for "Divergent regulation of STING-mediated innate immune responses by CBP and p300"

**[#]Corresponding to**

Mustafa Tekin, M.D., 1501 NW 10th Ave, BRB-610 (M860), Miami, FL 33136

Gaofeng Wang, Ph.D., 1501 NW 10th Ave, BRB-608 (M860), Miami, FL 33136

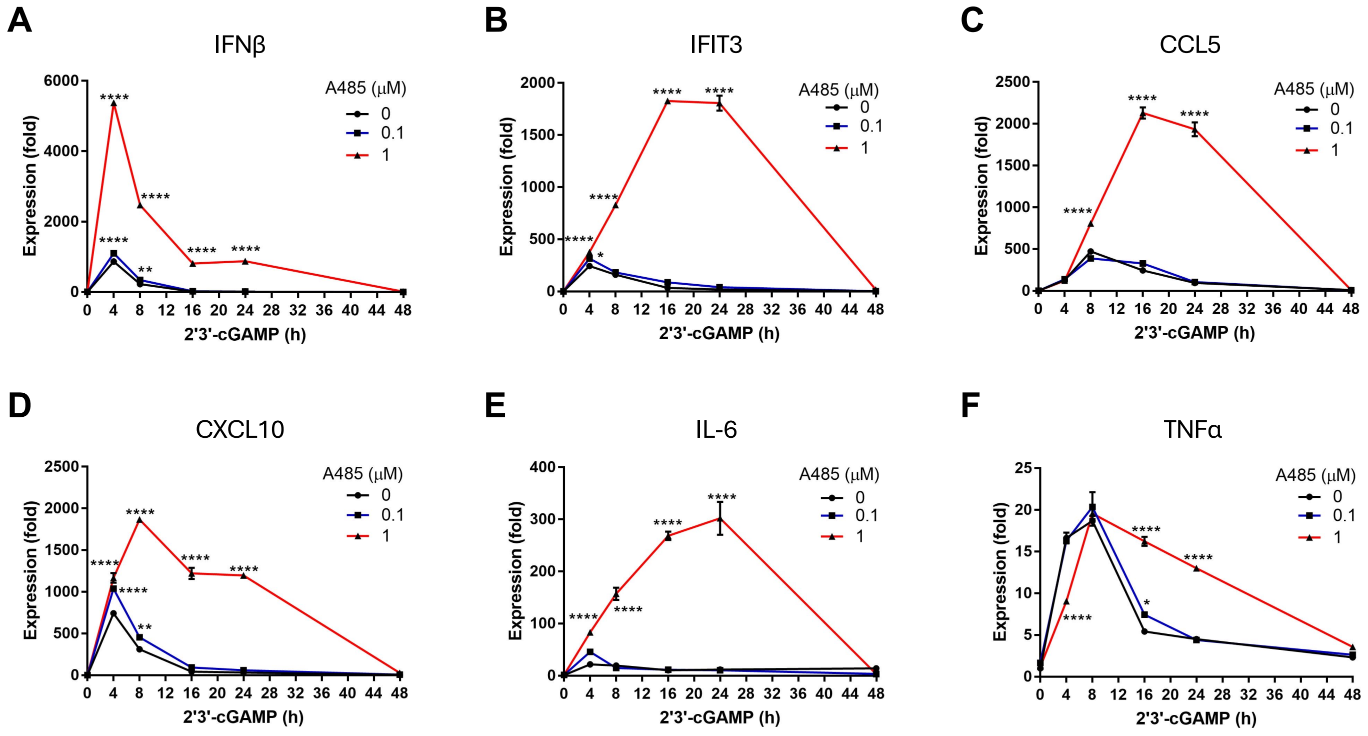


**Figure S1. p300i A485 enhances the cGAMP-induced transcription of innate immune genes in Raw264.7 cells.** (**A-F**) Pretreatment with A485 (0-1 μM) for 2 h simultaneously at the beginning of experiments dose-dependently enhances the transcription of innate immune responsive genes, including IFNβ, IFIT3, CCL5, CXCL10, IL-6 and TNFα induced by cGAMP (2 μM) in Raw264.7 cells. A485 alone has no effect on the transcription of these genes. * $P<0.05$, ** $P<0.01$, **** $P<0.0001$.

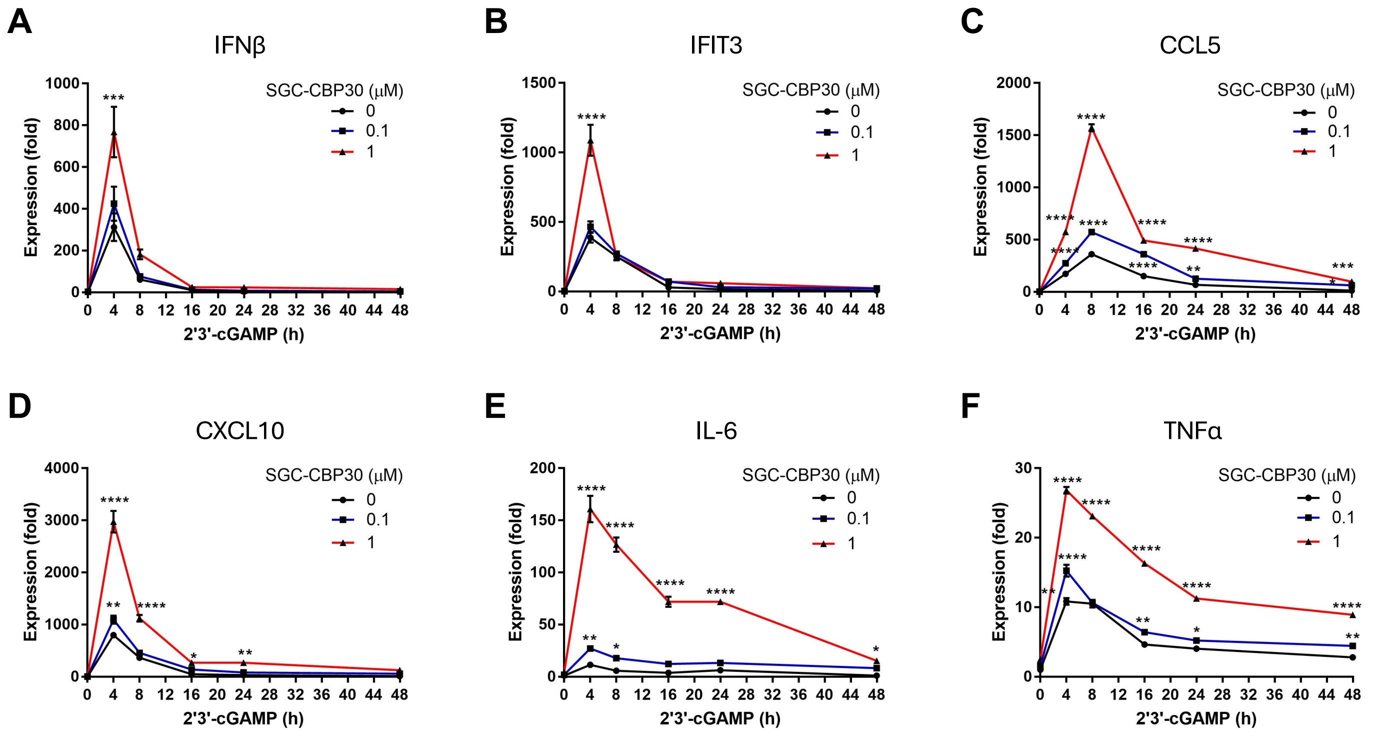


**Figure S2. p300i SGC-CBP30 enhances the cGAMP-induced transcription of innate immune genes in Raw264.7 cells.** (**A-F**) Pretreatment with SGC-CBP30 (0-1 μM) for 2 h simultaneously at the beginning of experiments dose-dependently enhances the transcription of innate immune responsive genes, including IFNβ, IFIT3, CCL5, CXCL10, IL-6 and TNFα induced by cGAMP (2 μM) in Raw264.7 cells. SGC-CBP30 alone has no effect on the transcription of these genes. * $P$<0.05, ** $P$<0.01, **** $P$<0.0001.

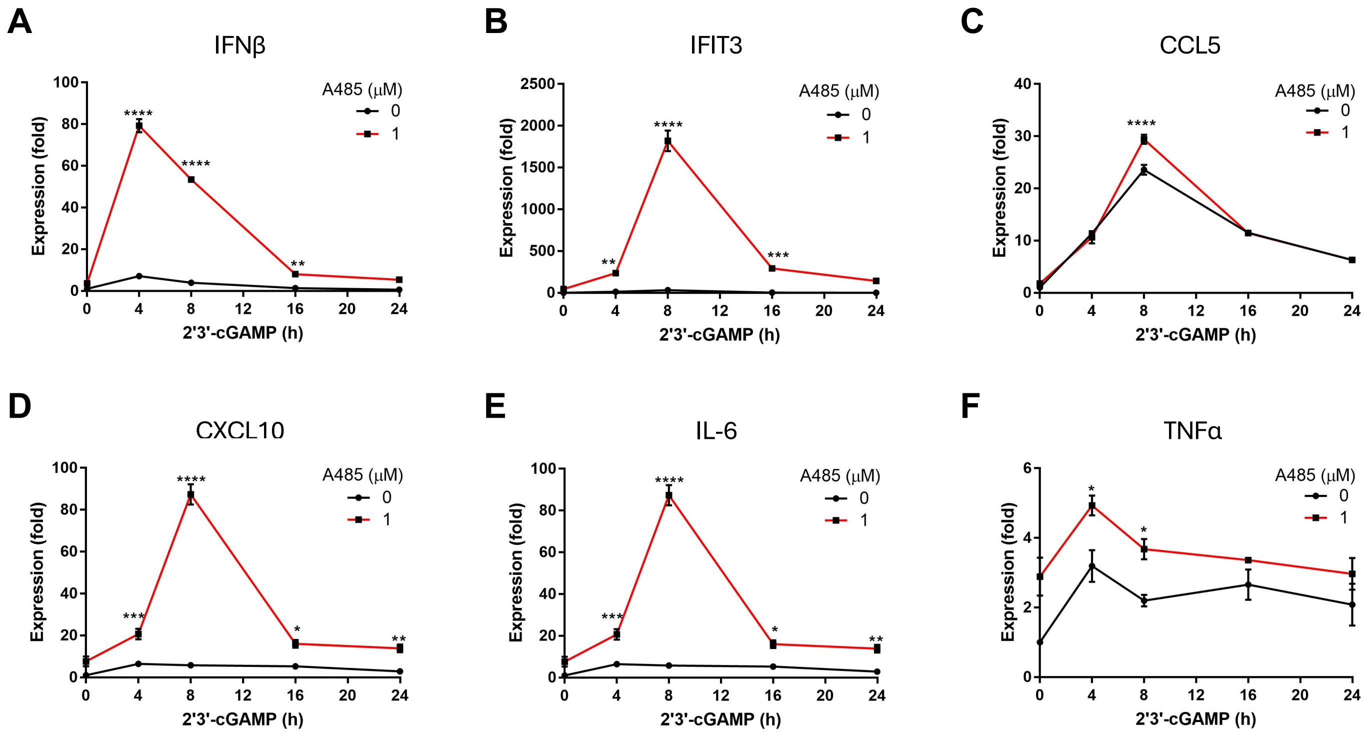


**Figure S3. p300i A485 enhances the cGAMP-induced transcription of innate immune genes in 4T1 cells.** (**A-F**) Pretreatment with A485 (1 μM) for 2 h simultaneously at the beginning of experiments enhances the transcription of innate immune responsive genes, including IFNβ, IFIT3, CCL5, CXCL10, IL-6 and TNFα induced by cGAMP (2 μM) in 4T1 tumor cells. A485 alone has no effect on the transcription of these genes. * $P<0.05$, ** $P<0.01$, **** $P<0.0001$.

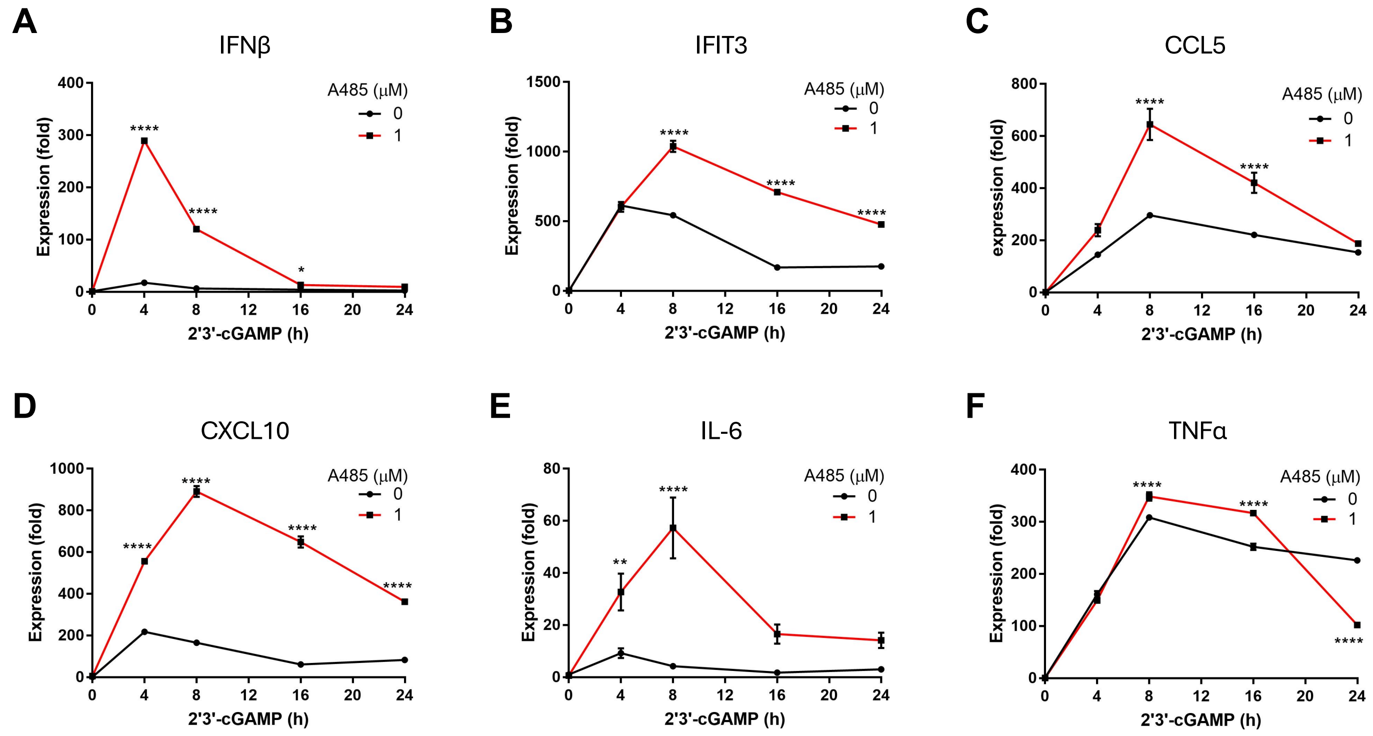


**Figure S4. p300i A485 enhances the cGAMP-induced transcription of innate immune genes in B16F10 cells.** (A-F) Pretreatment with A485 (1 μM) for 2 h simultaneously at the beginning of experiments enhances the transcription of innate immune responsive genes, including IFNβ, IFIT3, CCL5, CXCL10, IL-6 and TNFα induced by cGAMP (2 μM) in B16F10 tumor cells. A485 alone has no effect on the transcription of these genes. * $P<0.05$, ** $P<0.01$, **** $P<0.0001$.

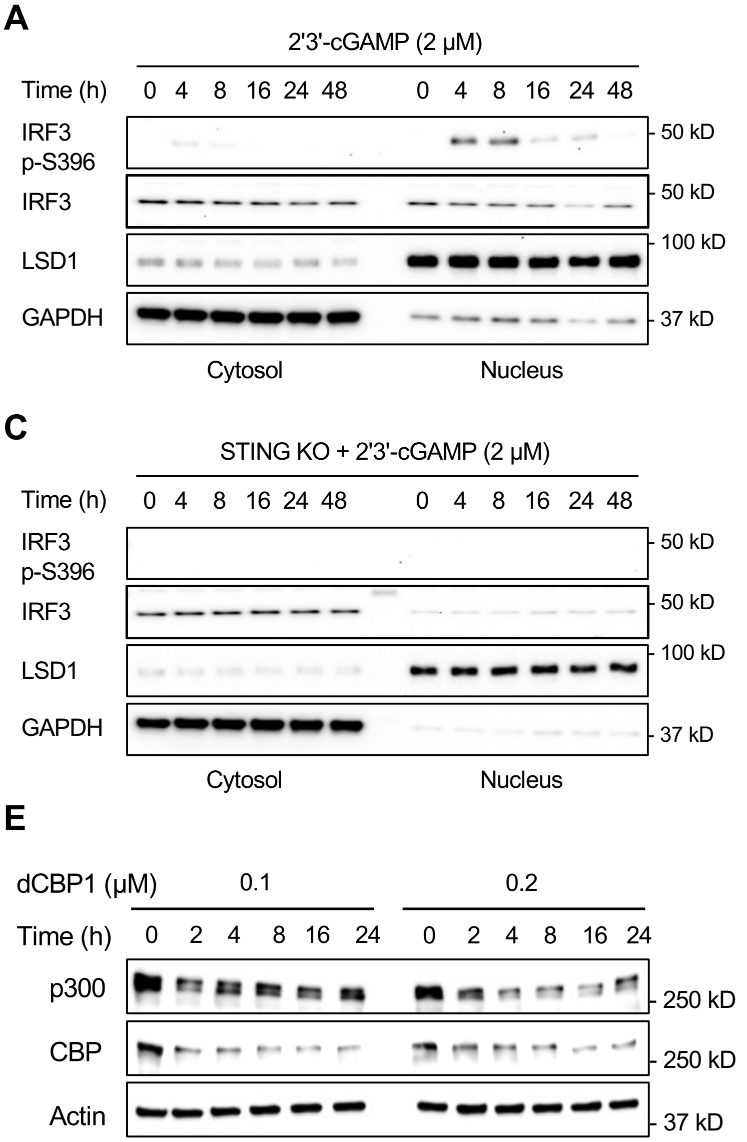


**Figure S5. TBK1 inhibitor and knockout of STING abolish the cGAMP-induced IRF3 phosphorylation.** (**A**) cGAMP (2 μM) induces IRF3 phosphorylation in Raw264.7 cells. (**B**) TBK1 inhibitor GSK8612 (1 μM) blocks the cGAMP-induced IRF3 phosphorylation. (**C**) Knockout of STING abolishes the cGAMP-induced IRF3 phosphorylation. (**D**) Validation of STING KO in different clones by immunoblot using actin as an inner control of whole cell lysates. LSD1: a nuclear marker; GAPDH: a cytosolic marker. (**E**) dCBP1 degrades both CBP and P300 with a slightly stronger effect on CBP in Raw264.7 cells. (**F**) No acetylation of TBK1 is detected by co-IP before or after cGAMP stimulation, with or without p300i A485 (1 μM) pretreatment in Raw264.7 cells

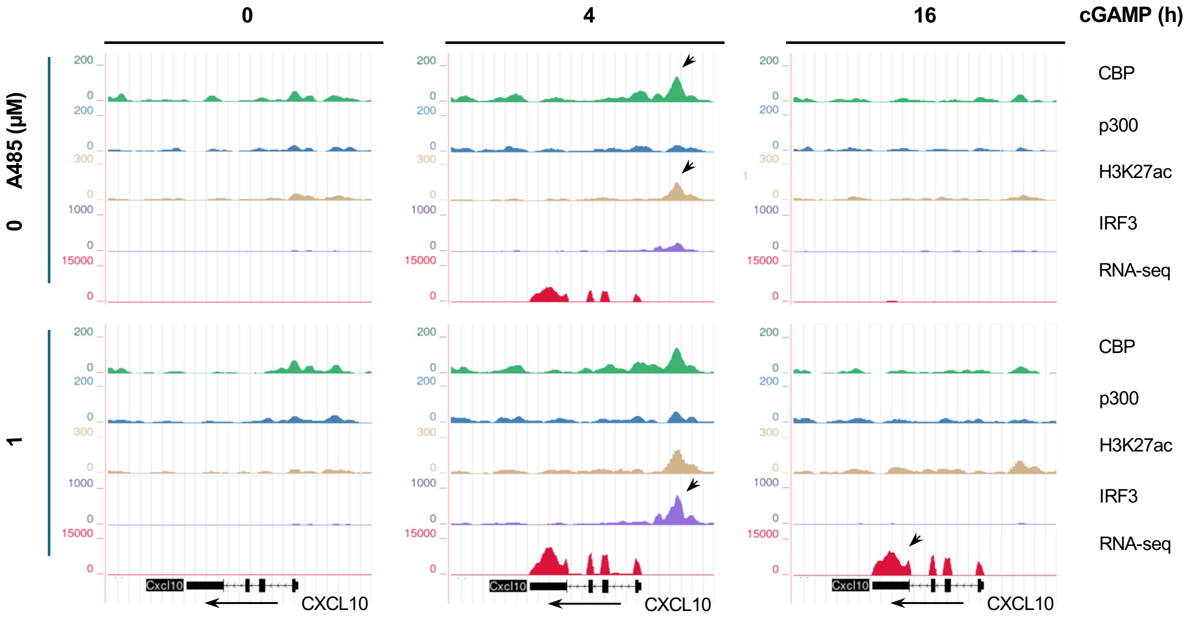


**Figure S6. p300i promotes the cGAMP-induced transcription by enhancing IRF3 chromatin recruitment.** Sequencing traces show that cGAMP induces the transcription of CXCL10, which correlates with the upregulated peaks of IRF3, H3K27ac (arrow) and CBP (arrow), but not p300 which remains largely unchanged, at the promoter region. p300i A485 increases and extends the transcription of CXCL10 (arrow), which correlates with the elevated IRF3 peak (arrow) at the promoter region.

**Table S1. Primers for murine innate immune genes.**

| Gene | Forward | Reverse |
|---|---|---|
| **ifnβ** | ATAAGCAGCTCCAGCTCCAA | CTGTCTGCTGGTGGAGTTCA |
| **cxcl10** | CCAAGTGCTGCCGTCATTTTC | GGCTCGCAGGGATGATTTCAA |
| **ccl5** | TTTGCCTACCTCTCCCTCG | CGACTGCAAGATTGGAGCACT |
| **ifit1** | GCAGAGAGTCAAGGCAGGTT | TTGTGCATCCCCAATGGGTT |
| **ifit3** | TTCCCAGCAGCACAGAAACA | AATGGCACTTCAGCTGTGGA |
| **il6** | AGTTGCCTTCTTGGGACTGATG | GGGAGTGGTATCCTCTGTGAAGTCT |
| **tnf** | CCAAATGGCCTCCCTCTCAT | TGGTGGTTTGCTACGACGTG |
| **gapdh** | AACGACCCCTTCATTGACCT | TGGAAGATGGTGATGGGCTT |
